# Therapeutic effects of a purified chitin-binding protein from *Moringa oleifera* seeds on irinotecan-induced intestinal mucositis in mice

**DOI:** 10.1101/2024.11.27.625542

**Authors:** Kayanny Queiroz Ferreira, Luana David do Carmo, Liviane Maria Alves Rabelo, Tiago Deivison Perreira Lopes, Deysi Viviana Tenazoa Wong, Aurilene Gomes Cajado, Renata Ferreira de Carvalho Leitão, Daniele Oliveira Bezerra Sousa, Nylane Maria Nunes de Alencar

## Abstract

Intestinal mucositis is a common side effect of irinotecan-based anticancer regimens, affecting approximately 85% of colorectal cancer (CRC) patients undergoing chemotherapy. Current treatment options are mainly palliative. Moringa oleifera Lamarck, native to Northeast India, is known for its nutritional and therapeutic properties. Our research group has demonstrated that MoCBP4 (11.78 kDa), a thermostable chitin-binding protein isolated from Moringa oleifera seeds, possesses potent antinociceptive, antifungal, wound-healing, and anti-inflammatory activity, both orally and intraperitoneally. Therefore, this study aimed to evaluate the protective effect of Mo-CBP4 in a model of irinotecan (CPT-11)-induced intestinal mucositis. This study was approved by the Animal Research Ethics Committee of UFC – CEPA (9892300120) and (7796300120). Male Swiss mice (25-30 g) were divided into 3 groups: Group 1 received saline solution (0.9%, i.p.) once a day for seven days; Group 2 received irinotecan (75 mg/kg, i.p.) once a day for four days; Group 3 was treated for 7 days with Mo-CBP4 at a dose of 10 mg/kg e.v., respectively, 30 minutes before CPT-11, which was administered for 4 days. During the seven days, weight loss, diarrhea scores, and survival were evaluated. On the seventh day, blood was collected for leukocyte count, followed by euthanasia for duodenum collection and evaluation of the following parameters: small intestine length, intestinal contractility, histopathological and morphometric alterations, MPO, GSH, MDA, NO, cytokines (IL-1β, IL-6, KC, TGF-β, and IL-10), and PCR (IL-33, IL-17, Claudin-2, Occludin, and ZO-1). We evaluated the cytotoxicity of MoCBP4 in normal and tumor cells for 24h and 72h using the Sulforhodamine B (SRB) assay and its interference with the effect of SN38 [2.5μM] (active metabolite of CPT-11) for 48h in murine colon cancer (MC38). We injected MC38 cells into male C57BL/6 mice (n=10; 25±2g) to induce a tumor amenable to treatment with CPT-11. We analyzed daily weight and palpable tumor growth with a digital caliper, and after sacrifice, we measured final tumor growth and MPO. Irinotecan (CPT-11) induced intestinal mucositis in mice, characterized by weight loss, diarrhea, increased mortality, leukopenia, decreased intestinal length, increased intestinal contractility, histopathological alterations (villous blunting, loss of crypt architecture, vacuolation, inflammatory cell infiltrate), and increased levels of MPO, IL-1β, IL-6, KC, TGF-β, IL-33, and Claudin 2. Increased MDA and decreased GSH levels were also observed. Treatment with Mo-CBP4 (10 mg/kg) significantly attenuated CPT-11-induced intestinal mucositis, improving diarrhea, increasing survival, reducing intestinal damage, and attenuating histopathological alterations. MoCBP4 was also able to decrease levels of MPO (35%), NO (48%), IL-1β (52%), IL-6 (98%), KC (88%), TGF-β (62%), IL-33, decrease MDA levels, and increase GSH. MoCBP4 did not exhibit cytotoxic activity in MC38, L929, SK-MEL, and B16F10 cell lines, and did not interfere with the cytotoxic effect of SN38. It also did not interfere with the antitumor effect of CPT-11 in a tumor transplant model. Therefore, Mo-CBP4 demonstrates important antidiarrheal, anti-inflammatory, and antioxidant activities that make it a promising therapeutic option for preventing and attenuating the severity of intestinal mucositis during CPT-11 chemotherapy treatment without interfering with the antitumor effect of irinotecan.

## Introduction

Colorectal cancer (CRC) is a global public health problem, with 1.9 million new cases and 930,000 estimated deaths in 2020. Incidence rates are higher in Australia/New Zealand and European regions, while they are lower in several African regions and South Asia (1). By 2040, there are projected to be 3.2 million new cases and 1.6 million deaths, mainly in countries with high or very high development indices. In Brazil, CRC ranks third in the list of cancer occurrences, being one of the types of cancer that has caused the most morbidity and mortality worldwide in recent years (2).

Irinotecan (CPT-11), a topoisomerase I inhibitor, is widely used in the treatment of various types of cancer, including CRC. CPT-11 is a prodrug that, after conversion by the hepatic carboxylesterase (CE) enzyme, produces its active metabolite, SN-38 (7-ethyl-10-hydroxycamptothecin). SN-38 is 1000 times more potent than CPT-11 and is responsible for the efficacy and toxicity of the drug. Prolonged exposure to SN-38 causes intense damage to the intestinal mucosa, leading to irinotecan toxicity (3,4).

Intestinal mucositis (IM) is a common side effect of irinotecan treatment, affecting about 85% of CRC patients(5). IM is characterized by inflammation, ulcerations, and erythema in the lining of the gastrointestinal (GI) tract. IM is associated with nausea, diarrhea, vomiting, cramps, anal pain, and abdominal distension. Despite being a widely discussed side effect, IM is still poorly treatable (5–7).

*Moringa oleifera* Lamarck, native to Northeast India, is known for its nutritional and therapeutic properties. *Moringa oleifera* seed is rich in polyphenols and has potent reducing power, free radical scavenging activity, and hydroxyl radical scavenging activity, making it a promising source of antioxidant bioactive compounds (8). MoCBP4 is a chitin-binding protein purified from Moringa oleifera seed (9,10). MoCBP4 has potent antinociceptive, antifungal, wound-healing, and anti-inflammatory activity, both orally and intraperitoneally. MoCBP4 is a basic protein with an isoelectric point (pI) of 10.55, a molecular weight of 11.78 kDa, and was isolated by Pereira *et al*. (2011). Lopes *et al*. (2020) deciphered its three-dimensional structure, finding similarities to a 2S albumin. MoCBP4 is stable at different pHs and temperatures, behaves as a glycoprotein with 2.85% carbohydrate content, and is resistant to proteolysis. The protein demonstrated the ability to inhibit leukocyte migration in models of cystitis and pancreatitis, as well as to regulate cytokines such as IL-1β, TNFα, and IL10 in a skin wound model. Mo-CBP4 has also proven effective in protecting cells from the harmful effects of ROS (reactive oxygen species), being stable and non-toxic to humans (9,11–13). However, the effect of Mo-CBP4 on chemotherapy-induced damage, on the drug itself, and on some tumor cell lines is still unknown. This study aims to investigate the potential of Mo-CBP4 in attenuating irinotecan-induced intestinal mucositis, evaluating its safety and efficacy in in vivo and in vitro models.

## Material and Methods

### MoCBP4 Isolation and purification

Isolation and purification of MoCBP4 was made in accord with previous studies (9,10). *Moringa oleifera* seeds were sourced from trees located on the Pici Campus of the Federal University of Ceará (UFC), Fortaleza, Brazil. A voucher specimen (No. EAC34591) was deposited in the Prisco Bezerra Herbarium, UFC. All proteins eluted from the seeds were named Mo-CBP (Mo: *Moringa oleifera* and CBP: chitin-binding protein), followed by a number in the order they were eluted. The MoCBP4 peak exhibited the highest yield compared to the other peaks, making it the chosen fraction for biochemical characterization published by Lopes *et al.* (2022).

### Animals and experimental

The animals were housed in appropriate polypropylene cages with ad libitum food and water, at room temperature (25 ± 3 °C) under a 12 h dark/12 h light cycle. The experimental protocols were conducted in accordance with current guidelines for the care of laboratory animals and ethical guidelines, reviewed and approved by the Animal Research Ethics Committee of UFC, Brazil, under protocol number - CEPA (9892300120) and (7796300120).

For the model of irinotecan (CPT-11)-induced intestinal mucositis, male Swiss mice (*Mus musculus*) (n=10; 25±2g) were used. In the tumor transplant experiments male C57BL/6 mice (n=10; 25±2g) with induced tumors were used.

### Irinotecan-induced intestinal mucositis model

The experimental model of intestinal mucositis used was initially described by Araki *et al*. (1993); Ikuno *et al*. (1995) and adapted in the Laboratory of Pharmacology of Inflammation and Cancer (LAFICA), Federal University of Ceará. This model consists of four administrations of irinotecan via intraperitoneal (i.p.) injection. After these four administrations, the animals present diarrhea and increased mortality. On the seventh day, the animals are euthanized, and the intestine is collected for the determination of general parameters of intestinal mucositis induction(14–17).

Briefly, mice were randomly assigned to receive either saline solution (3.5 mL/kg, i.p.) representing the vehicle group, or MoCBP4 (10mg/kg) 1 day before the administration of irinotecan (75 mg/kg i.p.; Trebyxan®; Bergamo, São Paulo, Brazil). From the first to the fourth day, the animals received irinotecan (75 mg/kg, i.p.) . Besides, from days 1 to 7, the animals were administered daily MoCBP4 (i.v.) or saline solution (3.5 mL/kg, i.p.) 1 h before irinotecan administration. the animals were assessed for diarrhea, blood leukocyte count, and body weight. Animals were euthanized with an overdose of ketamine/xylazine (>100/10 mg/kg, s.c.) for the collection of duodenal samples. These samples were then used for morphometric analysis, myeloperoxidase assay, oxidative stress assays, cytokine level determination, and Claudin-2 and IL-33 mRNA expression analysis.

#### Diarrhea Assessment

The severity of diarrhea was defined as described by Kurita *et al*. (2000) as follows: low: 0 (normal), normal stool; 1 (slightly), slightly wet stool; 2 (moderate), wet and formless stool with moderate perianal staining of the coat; and 3 (strong), a watery stool with strong perianal staining of the coat. Analysis was performed by an investigator blinded to the treatment.

#### Total Leukocyte Count

On the seventh day, blood was collected from all animals to assess the effect of irinotecan on leukopenia. Animals were anesthetized, and blood samples were collected from the ocular artery using a heparinized microcapillary tube. A 20 μL aliquot of blood was diluted in 380 μL of Turk’s solution and counted using a Neubauer chamber. The results were expressed as the total number of leukocytes x 103/mL of blood.

#### Histopathological analysis and intestinal morphometry

Duodenal segments were preserved in 10% (v/v) neutral buffered formalin, dehydrated, and embedded in paraffin. Sections of 5 μm thickness were stained with hematoxylin and eosin (H&E) for subsequent light microscopy examination. Mucosal injury was assessed using a modified histopathological scoring system adapted from Macpherson & Pfeiffer (1978). The length of the intestinal villi and depth of the crypts were measured using ImageJ software version 1.4 (NIH– National Institutes of Health, Bethesda, MD, USA). Five and 10 villi and crypts were measured per section, and 5-8 sections were analyzed per group.

### Myeloperoxidase (MPO) activity dosage

To assess the presence of neutrophils in the duodenal tissue samples, myeloperoxidase (MPO) activity was measured. MPO is an enzyme abundantly found in neutrophil granules, making it a reliable marker for neutrophil infiltration. On the day of the assay, the samples were homogenized in a buffer solution containing NaCl (100 mM), EDTA (15 mM), and NaPO4 (20 mM), followed by centrifugation. The pellet was resuspended in a buffer solution containing NaPO4 (50 mM) and hexadecyl-trimethyl-ammonium bromide, and centrifuged again. The supernatant was then used for the MPO assay. The assay was performed using tetramethylbenzidine (1.6 mM) and hydrogen peroxide (0.5 mM) as substrates. The enzymatic reaction was stopped by adding 50 µl of H2SO4 (1M) to each well. MPO activity was determined by measuring the absorbance at 450 nm using a microplate reader. The results were expressed as MPO activity per milligram of tissue.

#### Cytokine Quantification TNF-a, IL-1b, IL-6, IL-10 and TGF-b

To assess the levels of TNF-α, IL-1β, IL-6, IL-10, and TGF-β in the samples, a commercially available enzyme-linked immunosorbent assay (ELISA) kit (R&D Systems) was employed. Samples collected on the 7th day were analyzed for the presence of these cytokines. Briefly, 96-well microtiter plates were incubated with primary antibodies (R&D Systems) diluted in PBS (1:1000). Subsequently, the plates were incubated with biotinylated monoclonal antibodies against TNF-α, IL-1β, IL-6, IL-10, and TGF-β (R&D Systems) diluted in 1% BSA (1:1000). After washing, HRP-streptavidin (R&D Systems) was added to each well. The colorimetric reaction was initiated by adding o-phenylenediamine (R&D Systems) and incubated in the dark. The enzymatic reaction was stopped with 100 µl of stop solution (2 N H2SO4). The absorbance was measured at 450 nm using a microplate reader. The cytokine levels were expressed as picograms of cytokine per milligram of tissue

#### Gene Expression Analysis

Total RNA was extracted from duodenum tissue samples using TRIzol reagent (Invitrogen, Carlsbad, CA, USA) according to the manufacturer’s instructions. Briefly, 1 mL of TRIzol reagent was added to each sample and homogenized using a TissueLyser LT (Qiagen, Hilden, Germany) for 2 minutes at 50 Hz. The samples were then incubated on ice for 2 minutes and homogenized again. This procedure was repeated twice to ensure complete homogenization.

RNA was extracted from the samples using the TRIzol reagent (Invitrogen, Carlsbad, CA, USA) according to the manufacturer’s instructions. Briefly, 1 mL of TRIzol reagent was added to each sample and homogenized using a TissueLyser LT (Qiagen, Hilden, Germany) for 2 minutes at 50 Hz. The samples were then incubated on ice for 2 minutes and homogenized again. This procedure was repeated twice to ensure complete homogenization.

RNA was quantified using a NanoDrop 2000 spectrophotometer (Thermo Scientific, Waltham, MA, USA). The integrity of the RNA was assessed using an Agilent 2100 Bioanalyzer (Agilent Technologies, Santa Clara, CA, USA).

cDNA was synthesized from 1 μg of total RNA using the High Capacity cDNA Reverse Transcription Kit (Applied Biosystems, Foster City, CA, USA) according to the manufacturer’s instructions. Real-time PCR was performed using the QuantStudio™ 3 Real-Time PCR System (Applied Biosystems, Foster City, CA, USA) with PowerUp SYBR Green Master Mix (Applied Biosystems, Foster City, CA, USA). The PCR reaction was performed using the following primers:

**Table 1.**
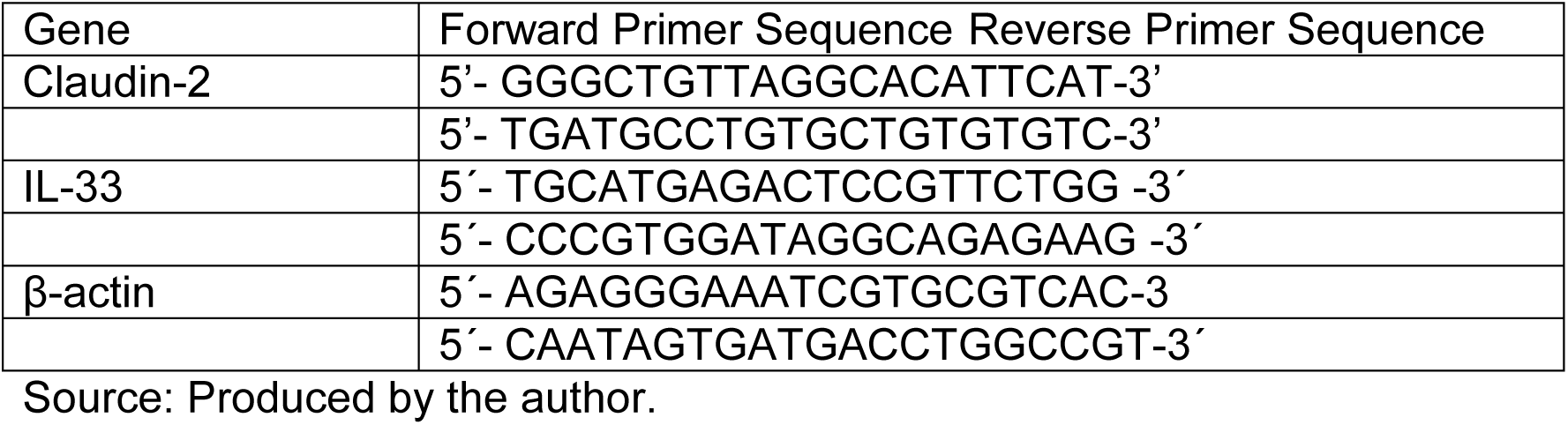
Primer sequences for real-time PCR.

The PCR reaction was performed using the following cycling conditions: 95°C for 10 minutes, followed by 40 cycles of 95°C for 15 seconds, 60°C for 1 minute, and 72°C for 1 minute. The relative gene expression was calculated using the 2-ΔΔCt method, with β-actin as the reference gene.

### Oxidative Stress Assessment

#### Malondialdehyde (MDA) Quantification

Malondialdehyde (MDA) is a product of the decomposition of polyunsaturated fatty acid hydroperoxides, formed during the oxidative process (5). The reaction involves 2-thiobarbituric acid with MDA, producing a red-colored compound, measured spectrophotometrically at 532 nm wavelength.

Duodenum tissue samples were homogenized in 0.05 M phosphate buffer (pH 7.4) at 10% (weight/volume) using a Polytron® homogenizer. Then, 250 μL of the homogenate were placed in a water bath at 37°C for 1 hour. To interrupt peroxidation, 400 μL of 35% perchloric acid were added to the samples, which were then centrifuged (14000 rpm, 15 minutes, 4°C). From the supernatant obtained, 600 μL were transferred to a microtube, to which 200 μL of 0.8% thiobarbituric acid were added. This mixture was placed in a water bath at 95°C for 30 minutes. After cooling, the samples were plated and the absorbance reading was performed in a microplate reader at a wavelength of 532 nm. A standard curve with known concentrations of tetramethoxypropane (TMP) was used to calibrate the method and the MDA concentration in the samples was calculated using the equation of the straight line for the standard curve. Results were expressed in nmol of MDA/g of tissue.

#### Reduced Glutathione (GSH) Levels

Reduced glutathione (GSH) levels were measured in the duodenum tissue samples. GSH is a water-soluble antioxidant recognized as the most important endogenous component of the pool of non-protein sulfhydryl groups (NPSH) in our body. To determine the concentration of GSH, the NP-SH content was analyzed using the technique described by Sedlak and Lindsay (1968), which is based on the reaction of 5,5-dithiobis(2-nitrobenzoic) acid (DTNB) with compounds of sulfhydryl, and consequent development of yellow coloration. DTNB reacts with GSH forming 2-nitro-5-thiobenzoic acid and oxidized glutathione (GSSG). The skin samples were homogenized at 10% (weight/volume) in Politron® with 0.02 M EDTA solution. Soon after, 60 μL of 10% trichloroacetic acid (TCA) were added to 40 μL of each sample in order to precipitate the proteins present in the biological material. The material was then centrifuged (5000 rpm, 15min, 4°C) and 60 μL of the obtained supernatant was plated. A standard curve with known GSH concentrations was used for method calibration. 102 μL of the reading solution (Tris-EDTA, DTNB 0.01 M) were added and the absorbance was immediately measured in a microplate reader at a wavelength of 412 nm. The GSH concentration in the samples was calculated using the straight-line equation for the standard curve and the results were expressed in μg of GSH/g of tissue(18).

#### NO conversion to NO2 and NO3 analysis

To determine nitrite (NO2-) levels, duodenum tissue samples were collected on the 7 day. The samples were weighed and homogenized using a Polytron® homogenizer in a cold solution of potassium chloride (1.15% KCl). After obtaining the homogenates of each sample (10% tissue), centrifugation was performed to obtain the supernatant (1500g; 15 minutes) then they followed the methodology Green *et al*, (1981) (19). The reading of the final purple color obtained by the reaction was performed at the absorbance of 540 nm and the results were expressed in μM of NO2-, using the standard nitrite curve as a reference to obtain the values (19,20).

### Cell Culture

Four adherent cell lines, obtained from the Rio de Janeiro Cell Bank, were employed in this study: murine fibroblast cell line (L929), colorectal carcinoma cell line (MC38), SK-Mel melanoma cell line (ATCC®), and metastatic melanoma cell line B16F10. Cell lines were kept in Dulbecco’s modified Eagle’s medium (DMEM: Gibco, Gaithersburg, USA) supplemented with 10% Fetal Bovine Serum (FBS) and 1% penicillin-streptomycin and maintained in a 5% CO2 atmosphere at 37°C using a Panasonic® incubator (model MCO 19AICUVH)

#### Cell Viability Assay

The SRB assay was employed to evaluate the cytotoxicity of MoCBP4 on the cell lines. Following trypsinization and counting, cells were seeded in 96-well plates and incubated. After the incubation period, the culture medium containing any dead cells was carefully removed to avoid disturbing the viable adherent cells. The remaining cells were fixed with 100 µL of 10% (w/v) trichloroacetic acid (TCA) for at least 1 h at 4°C. After fixation, the cells were washed to remove any residual acid. Subsequently, 100 µL of 0.4% (w/v) sulforhodamine B (SRB) in 1% acetic acid was added, and the plates were incubated in a 5% CO2 atmosphere at 37°C for 30 min. The SRB dye binds to cellular amino acid residues. Following incubation, excess dye was removed by washing three times with 1% acetic acid. The bound SRB was then solubilized in 200 µL of 10 mM Tris-base buffer at 4°C and homogenized on a plate shaker for at least 5 minutes at room temperature. The absorbance was measured at 570 nm using a plate reader (Fisher Scientific, model Multiskan FC). The SRB results were expressed as a percentage of cell viability.

### Tumor transplant Assay

Mice were inoculated intramuscularly (i.m.) with 5x10⁵ MC-38 cells suspended in 50 µL of DMEM culture medium. Tumor growth was monitored daily using a digital caliper, and body weight was recorded to assess tumor progression. Thirty days post-inoculation, when tumors became palpable, mice were randomized into three treatment groups. Treatment continued for 7 days, after which the animals were euthanized. Duodenal tissue was collected for myeloperoxidase (MPO) assay, and tumor growth was assessed following tumor resection.

### Statistical analysis

Data are presented as mean ± standard error of the mean (SEM) for normally distributed data or median (minimum-maximum) for non-parametric data. Statistical analyses were performed using GraphPad Prism version 8.0. *In vitro* data (SRB assay) were analyzed by one-way ANOVA followed by Dunnett’s test. *In vivo* data were analyzed by one-way ANOVA followed by Bonferroni’s test for parametric data or Kruskal-Wallis test followed by Dunn’s test for non-parametric data. Survival analysis was performed using the Log-rank test. Differences were considered statistically significant at p < 0.05.

## RESULTS

### MoCBP4 ameliorates irinotecan-induced intestinal mucositis and its associated complications

Irinotecan administration resulted in a significant decrease in survival (58%) compared to the control group (100% survival). Treatment with MoCBP4 (10 mg/kg, i.v.) significantly improved survival rates (90%) compared to the irinotecan group (Fig. 1A). Consistent with the improved survival, MoCBP4 also ameliorated irinotecan-induced diarrhea. Animals treated with irinotecan alone exhibited diarrhea scores of 2 (0–3), while MoCBP4 treatment significantly reduced diarrhea severity (Fig. 1B). Furthermore, irinotecan administration caused a significant reduction (84.34%) in total leukocyte counts compared to the control group. MoCBP4 treatment resulted in a 59.64% reduction in leukocytes (Fig. 1C) compared to the control, representing a 24.7% improvement compared to the irinotecan group. MoCBP4 treatment did not alter the weight loss associated with irinotecan.

**Figure 1.**
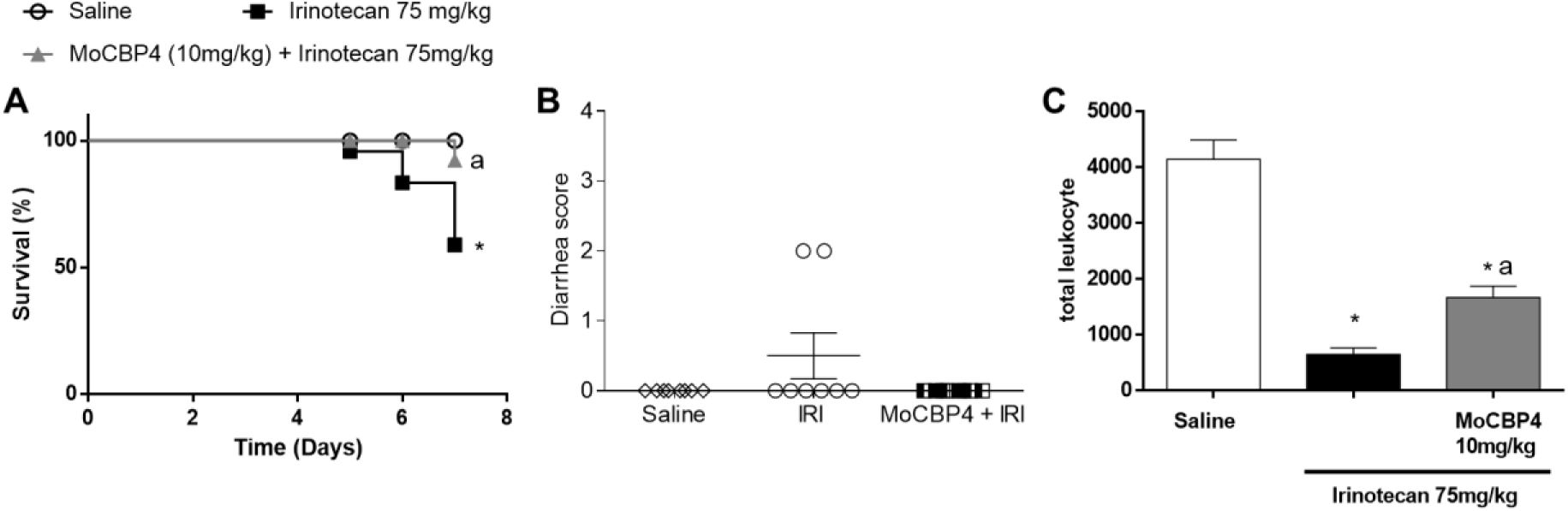
Effects of co-administration of MoCBP4 (10 mg/kg; i.v) on irinotecan-induced intestinal mucositis on **A** survival rate, **B** diarrhea, and **C** total leukocyte in mice with intestinal irinotecan-induced mucositis (saline: n = 8, MoCBP4: n = 10; the number of animals was doubled in the irinotecan-treated group (n = 20) to obtain an adequate number of mice for further analyses). **\***p < 0.05 compared to the saline group and **a** p < 0.05 compared to the saline + irinotecan-treated group (Mantel-Cox log-rank test; Kruskal–Wallis with Dunn’s test, and ANOVA with Bonferroni’s test, respectively.

### MoCBP4 reduces the histological damage and intestinal shortening induced by irinotecans

Irinotecan administration induced significant histological damage in the duodenum, characterized by shortened villi, loss of crypt architecture, vacuolization, and infiltration of inflammatory cells in the lamina propria and mucosa. The control group (saline) exhibited normal duodenal morphology. Irinotecan treatment significantly reduced small intestine length (25.51%) compared to the control group. MoCBP4 (10 mg/kg, i.v.) treatment preserved intestinal length, demonstrating a 36% increase compared to the irinotecan group, with no significant difference compared to the saline group (Fig. 2A). MoCBP4 treatment also significantly ameliorated the histological damage induced by irinotecan in the duodenum. Histopathological scores were significantly lower in the MoCBP4-treated group [1 (0–1)] compared to the irinotecan group [3 (3–4)], with the saline group exhibiting a score of 0 (0–0) (Fig. 2B). Furthermore, irinotecan treatment induced a significant inflammatory response in animals, characterized by intestinal shortening (Fig. 2C). This irinotecan-induced intestinal shortening was attenuated by MoCBP4 treatment, significantly improving the villus/crypt ratio (3.2±0.2) compared to the irinotecan group (2.5±0.1), as shown in Figure 2D.

**Figure 2.**
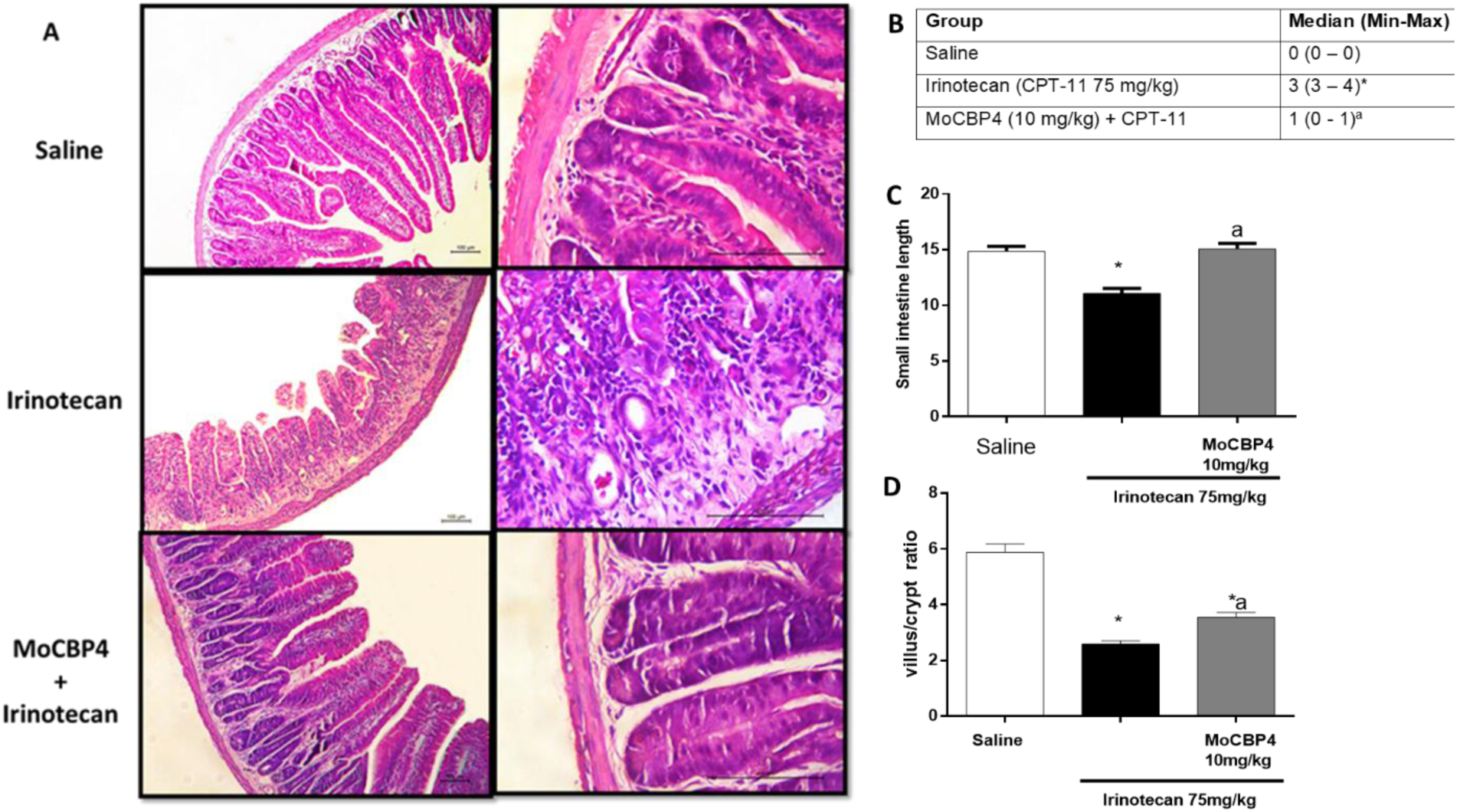
Histology of duodenum. **A** Representative histology of duodenum with hematoxylin and eosin stain in mice submitted to treatment with saline, saline + irinotecan, or MoCBP4 10 mg/kg (i.v.) + Irinotecan (×100 and ×400 magnifications). Scale bar: 100 µm. **B** Histological scores of the animals submitted to treatment with saline, MoCBP4 10 mg/kg + irinotecan (i.v.) (5–6 tissue sections were analyzed for each experimental group; saline: n = 5, irinotecan: n = 5, MoCBP4: n = 6). **C** Small intestine length of mice with irinotecan-induced intestinal mucositis. **D** villus/crypt ratio of duodenum mucosa. The values are expressed as the means ± SEM (parametric data) or median (minimum-maximum) for nonparametric data. **\***p < 0.05 compared to the saline group and **a** p < 0.05 compared to the irinotecan-treated group. Parametric and nonparametric data were analyzed using ANOVA followed by Bonferroni’s post hoc testing or Kruskal-Wallis/Dunn’s test.

### MoCBP4 attenuates the inflammatory response to mucositis

Neutrophil infiltration in the duodenum, as measured indirectly by myeloperoxidase (MPO) activity, was significantly increased in the irinotecan group (9762±6174) compared to the saline group (1134±423.9) (Fig. 3A). MoCBP4 (10 mg/kg, i.v.) treatment significantly reduced MPO activity (8203±2584) in the irinotecan-induced mucositis group. As expected, irinotecan treatment increased pro-inflammatory cytokine levels (IL-1β, IL-6, KC, and TGF-β) compared to the control group. MoCBP4 treatment significantly attenuated these increases, reducing IL-1β, IL-6, KC, and TGF-β levels by 52%, 98%, 88%, and 62%, respectively, compared to the irinotecan group (Fig. 3B-E). No significant differences in IL-10 levels were observed between the groups (Fig. 3F).

**Figure 3.**
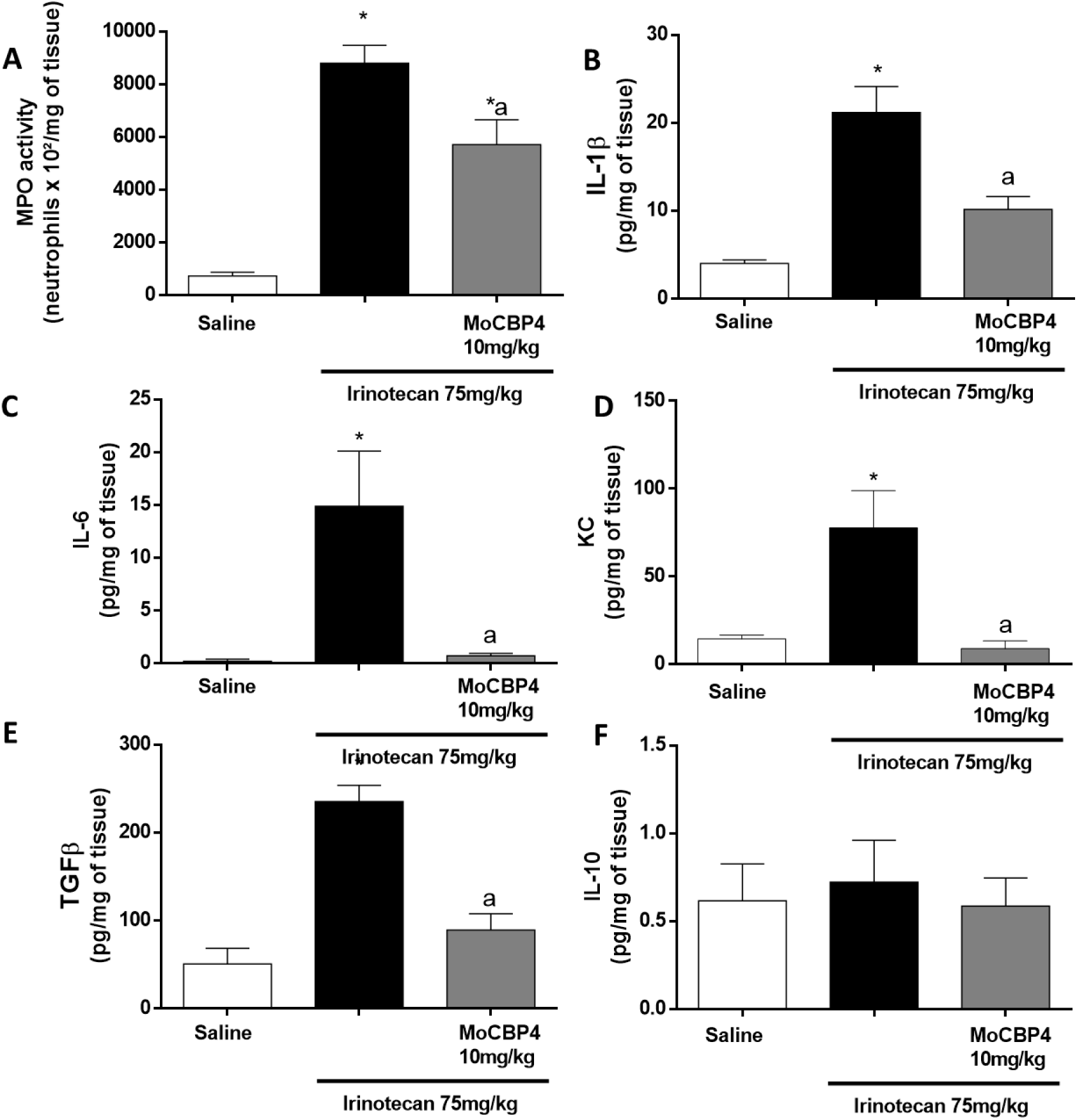
Modulation the Inflammatory Response After Mucositis Induction. MoCBP4 treatment reduced **A** MPO, **B** IL-1β, **C** IL-6, **D** KC, **E** TGF-β and **F** IL-10 activities in mice with irinotecan-induced intestinal mucositis. The results are expressed as the mean ± SEM. **\***p < 0.05 compared to the saline group and a p < 0.05 compared to the saline + irinotecan-treated group (Saline: n = 8, irinotecan: n = 8, MoCBP4: n = 8 animals per group; ANOVA with Bonferroni’s test).

### IL-33 and Claudins-2 are expressed during intestinal mucositis

Duodenal tissue samples were analyzed for gene expression of inflammatory markers. Irinotecan treatment significantly increased IL-33 gene expression (Vehicle: 1.84 ± 0.37 vs. CPT-11: 11.91 ± 0.75) compared to the control group, and MoCBP4 treatment reversed this increase (p<0.05) (Fig. 4A). Irinotecan treatment also increased Claudin-2 expression (Vehicle: 2.05 ± 0.30 vs. CPT-11: 6.94 ± 0.91). MoCBP4 treatment did not reverse the irinotecan-induced increase in Claudin-2 expression (MoCBP4: 7.80 ± 1.10) (Fig. 4B).

**Figure 4.**
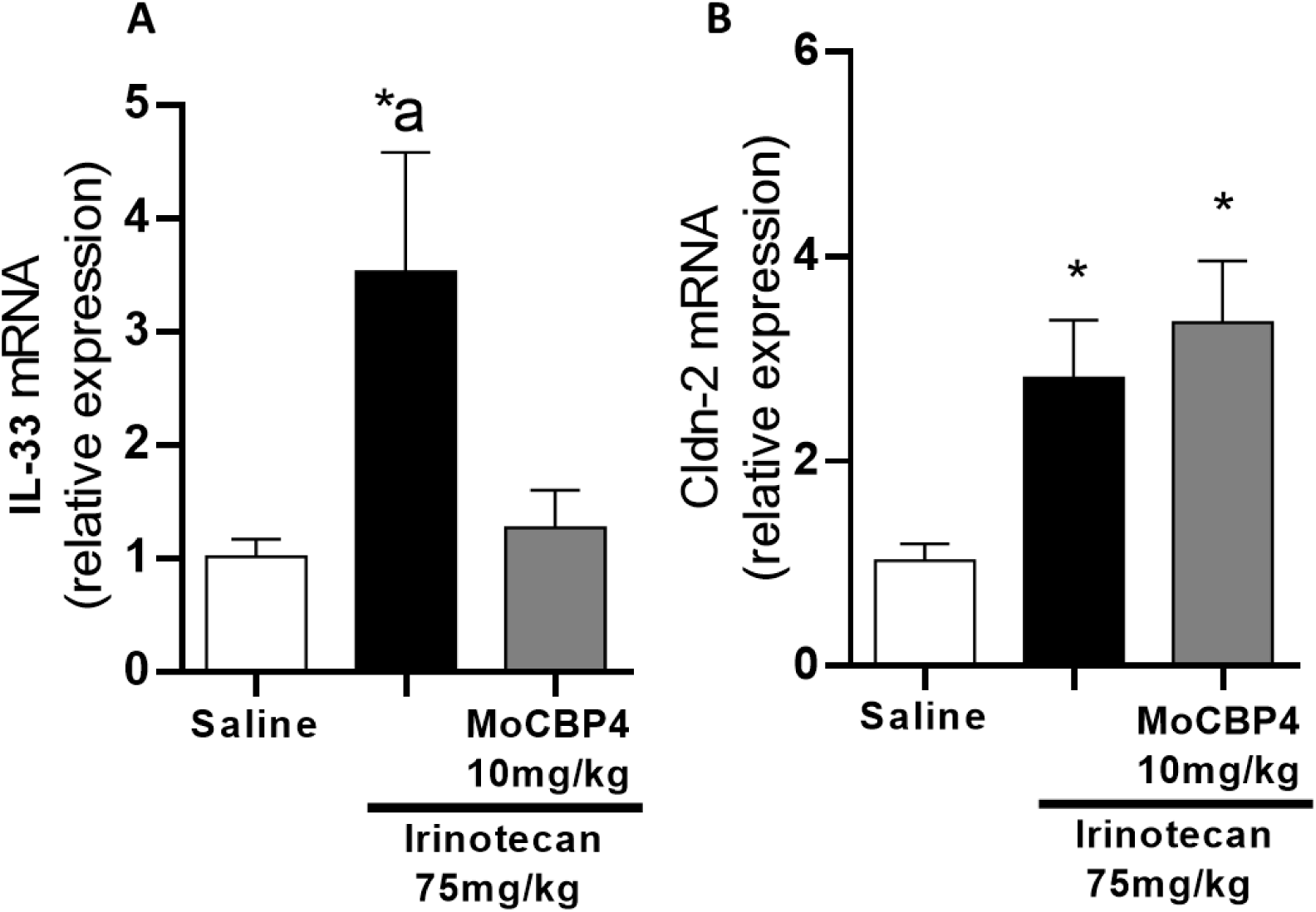
Gene expression of inflammatory mediators after mucositis induction. MoCBP4 treatment reduced A IL-33 mRNA relative expression. MoCBP4 treatment did not reverse the irinotecan-induced increase in Claudin-2 expression. The results are expressed as the mean ± SEM. *p < 0.05 compared to the saline group and a p < 0.05 compared to the saline + irinotecan-treated group (Saline: n = 8, irinotecan: n = 8, MoCBP4: n = 8 animals per group; ANOVA with Bonferroni’s test).

### MoCBP4 decreases Oxidative Stress

Assessment of oxidative stress markers in duodenal tissue revealed that irinotecan treatment alone did not alter MDA and GSH levels compared to the control group but increased NO levels by 104%. MoCBP4 (10 mg/kg, i.v.) treatment significantly reduced MDA and NO levels by 85.3% and 47.4%, respectively, and increased GSH levels by 865.75% compared to the irinotecan group (Fig. 5 A-C).

**Figure 5.**
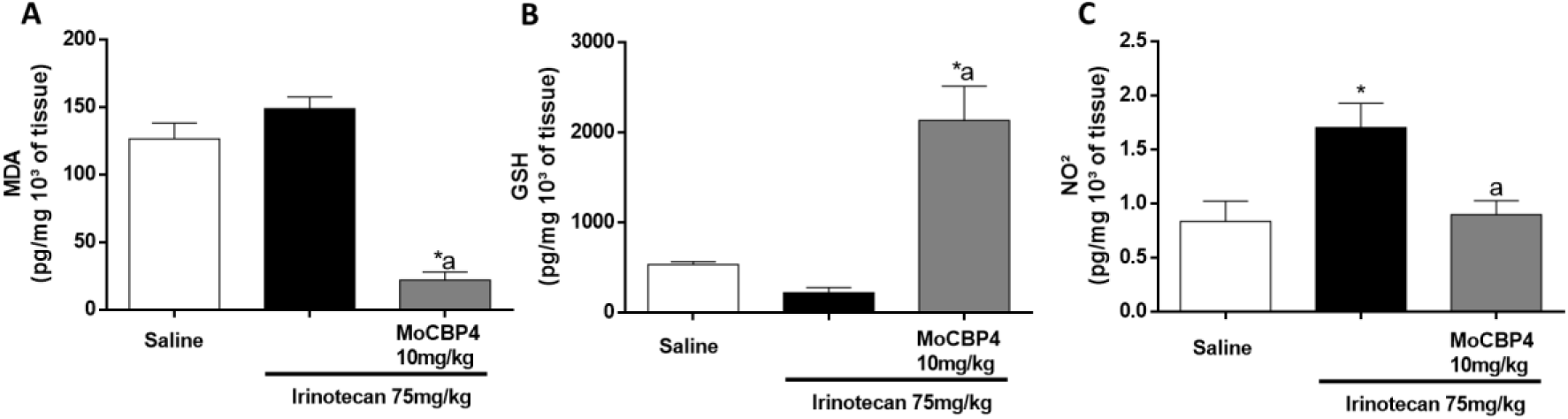
Oxidative stress was assessed by measuring A MDA, B GSH and C NO2/NO3 of tissue. The results (picogram/mg 10³ of tissue) were expressed as the mean ± standard error of the mean. ANOVA and Tukey’s posttest were used for comparisons between means. *p<0.05 represents a statistically significant difference in relation to the Saline group, a p<0.05 in relation to irinotecan (N=5-6 animals/group).

### MoCBP4 Maintains SN38 Cytotoxicity *In Vitro*

*In vitro* cytotoxicity of MoCBP4 was assessed in L929, SK-MEL, MC38, and B16F10 cell lines at 24 and 72 hours . Cell densities were adjusted according to their respective doubling times to ensure consistent treatment conditions. MoCBP4 did not inhibit cell growth in any of the tested cell lines (Tab. 1) Following concentration-response curves, four non-cytotoxic concentrations of MoCBP4 (6, 3, 1.5, and 0.7 µg/mL) were selected for further analysis with SN38 in MC38 cells, as these concentrations maintained cell viability above 90% after 48 hours. A concentration-response curve for SN38 was generated (0.3 - 40 µM), and the IC40 value (2.5 µM) was selected for co-treatment experiments. Co-treatment of MC38 cells with MoCBP4 and SN38 for 48 hours revealed that MoCBP4 did not negatively interfere with the cytotoxic effects of SN38 (Fig. 6). In vivo studies were subsequently conducted to further evaluate the antitumor effects of irinotecan with and without MoCBP4.

**Table 1.** Cytotoxic activity using the SRB assay in different cell lines.

| Cell line | Time | IC<sub>50</sub> | Time | IC<sub>50</sub> |
| --- | --- | --- | --- | --- |
| L929 | 24h | > 100μg/mL | 72h | > 100μg/mL |
| MC38 | 24h | > 100μg/mL | 72h | > 100μg/mL |
| SK-MEL | 24h | > 100μg/mL | 72h | > 100μg/mL |
| B16F10 | 24h | NT | 72h | > 100μg/mL |

The results are representing the inhibitory concentration for 50% of the cells (IC<sub>50</sub>) and 95% range. Three independent experiments were performed in duplicate. NT represents Not Tested.

**Figure 6.**
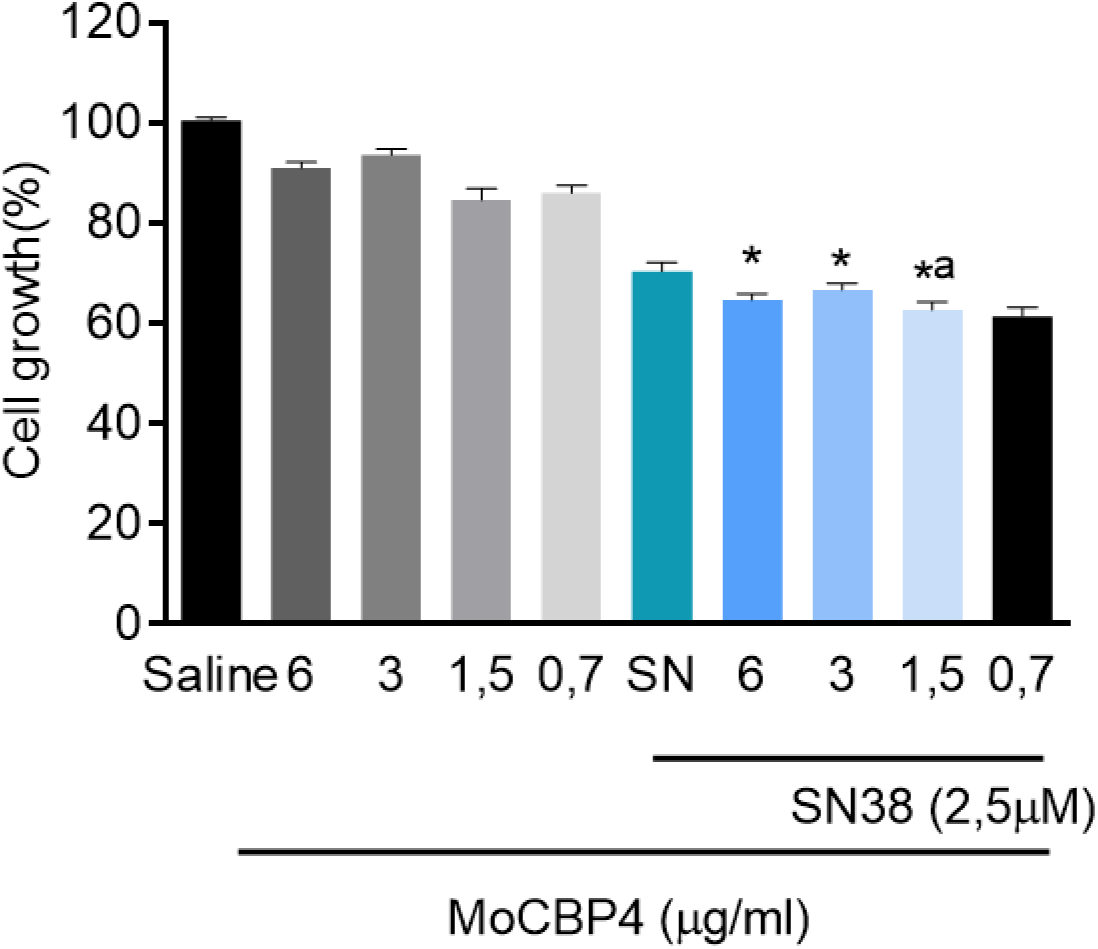
The cytotoxic activity of MoCBP4 in combination with SN38 was assessed in MC38 cells using the SRB assay. Cells were seeded in 96-well plates (2 x 10⁴ cells/mL) and incubated with saline (vehicle) or MoCBP4 (6, 3, 1.5, and 0.7 µg/mL) for 30 minutes prior to SN38 (2.5 µM) addition. After 48 hours, cells were incubated with 0.4% SRB solution for 1 hour and absorbance was measured at 570 nm. Data are presented as mean percentage of cell viability ± SEM from three independent experiments performed in duplicate. *p < 0.05 compared to the corresponding concentration without SN38. ᵃp < 0.05 compared to SN38 alone. One-way ANOVA followed by Tukey’s test was used for statistical analysis.

### *In Vivo* Antitumor Effects of Irinotecan are Maintained with MoCBP4 Co-treatment

Tumor size and body weight were monitored daily. Following treatment, tumors were excised and measured, and duodenal tissue was collected for MPO assay. Irinotecan treatment resulted in weight loss in groups II and III, while MoCBP4 did not significantly affect this weight loss compared to irinotecan alone (Fig.7A). Although tumor size increased in all groups, irinotecan significantly limited tumor growth. MoCBP4 did not interfere with irinotecan’s antitumor effect, as demonstrated by similar tumor growth rates in both irinotecan-treated groups (Fig. 7 B-C). MPO activity in duodenal tissue was significantly increased in the irinotecan group, confirming the drug’s systemic effect on intestinal tissue. While MoCBP4 + irinotecan treatment also increased MPO activity, the increase was less pronounced than with irinotecan alone (Fig. 7 D).

**Figure 7.**
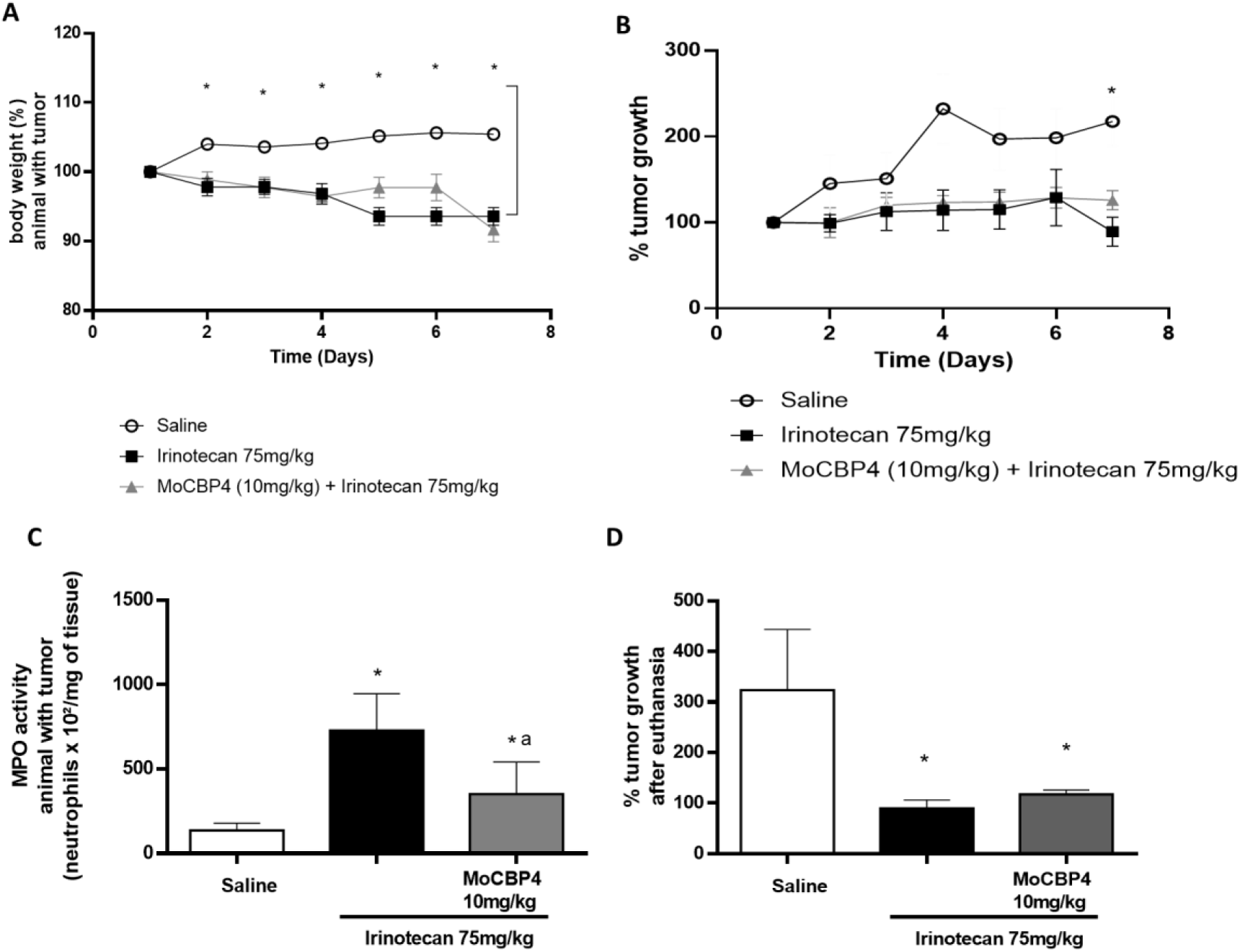
Effect of MoCBP4 on irinotecan treatment in a MC38 murine tumor model. C57BL/6 mice received MC38 cell transplants and were divided into three groups (n=10): (I) Control (saline, 5 mL/kg, i.p.); (II) Irinotecan (CPT-11, 75 mg/kg, i.p., 4 days); and (III) MoCBP4 (10 mg/kg, i.p., 7 days) + CPT-11 (75 mg/kg, i.p., 4 days). The graphs show the percentage change in body weight (A), the percentage of tumor growth during treatment (B), tumor growth compared to baseline (C), and duodenal myeloperoxidase (MPO) activity on day 7 (D). *p<0.05 vs. saline group; **a** p<0.05 vs. CPT-11 group (ANOVA, Bonferroni test).

### Discussão

*Moringa oleifera*, known for its therapeutic properties in various conditions, including cancer, diabetes, and inflammatory processes is the source of MoCBP4, a chitin-binding lectin with pharmacological potential (21–24). Plant proteins, as demonstrated in previous studies have shown promise in the treatment of different pathologies. In this study, we demonstrated, for the first time, the protective effect of MoCBP4 in irinotecan-induced intestinal mucositis, without interfering with the drug’s antitumor activity. MoCBP4 exerted antidiarrheal, anti-inflammatory, and antioxidant effects, highlighting its potential as a novel therapy for this debilitating condition (23,25).

The deleterious effects of chemotherapy, often resulting from the low selectivity of antineoplastic drugs and multidrug resistance, can lead to severe complications, including death (26–28). The search for strategies that mitigate these effects without compromising treatment efficacy is essential. Intestinal mucositis, a significant adverse effect in cancer patients, is particularly debilitating in the treatment of colorectal cancer (29–32). The irinotecan-induced mucositis model, adapted from Araki *et al*. (1993) and Ikuno *et al*. (1995), reproduces leukopenia, diarrhea, and alterations in the intestinal mucosa, corroborating previous findings (16,17,33). MoCBP4 significantly attenuated irinotecan-induced leukopenia, diarrhea, and mortality, increasing animal survival and demonstrating a protective effect on the intestinal mucosa. Irinotecan-induced diarrhea is associated with damage to the intestinal architecture, such as shortening of the intestine and impaired nutrient absorption (17). The results demonstrate that MoCBP4 prevents irinotecan-induced intestinal shortening, similar to that observed with other protective agents (34) and exerts an antidiarrheal effect, although this has not been previously reported for *Moringa oleifera* seed extracts (35). However, MoCBP4 did not interfere with irinotecan-induced weight loss.

The healing potential of MoCBP4, previously demonstrated in an excisional wound model (9), may contribute to the restoration of intestinal morphology and the consequent antidiarrheal effect observed in this study. MoCBP4 promoted epidermal reconstruction and proliferation of cells involved in healing, with complete wound closure in 12 days. The modulation of cytokines such as TNF-α, IL-1β, and IL-10, observed in the study by *Lopes et al*. (2022), suggests the activation of tissue repair mechanisms. Myeloperoxidase (MPO), a marker of neutrophil presence in inflamed tissues (36,37), was used to assess neutrophil infiltration in mucositis. Neutrophils, essential in the immune response to microorganisms, also contribute to tissue damage associated with chemotherapy (38,39). MoCBP4 reduced irinotecan-induced neutrophil migration, corroborating previous studies with other inflammatory models (13) and with lectins from different plant species(40–42). MoCBP4 also modulated the levels of pro-inflammatory cytokines (IL-1β, IL-6, KC, TGF-β), with significant reductions in irinotecan-treated animals, restoring IL-6, KC, and TGF-β levels to those of the control group. However, there was no change in IL-10 levels.

IL-33 plays a role in exacerbating intestinal mucositis by increasing neutrophil migration and causing mucosal damage (43) MoCBP4 reduced the irinotecan-increased levels of IL-33, suggesting a possible blockade of the IL-33/ST2 pathway, contributing to the observed protective effect. The increased expression of claudin-2 observed in irinotecan-treated animals may be related to increased IL-6 via STAT3 (44). Although MoCBP4 reduced IL-6 levels, there was no reduction in claudin-2 expression. This result may indicate a homeostatic response to repair the damaged intestinal epithelium (45) related to the suppression of tumor invasion (46). Further studies are needed to clarify the role of claudin-2 in mucositis and cancer.

Intestinal permeability, beyond the influence of claudins, is affected by oxidative damage, particularly from ROS and nitric oxide, contributing to irinotecan-induced diarrhea (16). Oxidative stress was assessed through measurement of reduced glutathione (GSH), malondialdehyde (MDA), and nitric oxide (NO) levels. MoCBP4 increased GSH levels and significantly reduced MDA and NO levels. ROS generation and the consequent oxidative stress, with DNA damage and activation of inflammatory pathways, constitute initial events in the pathogenesis of intestinal mucositis. Demonstrated the ability of MoCBP4 to reduce oxidative stress in a model of acute pancreatitis by increasing GSH levels, reducing MDA and nitrite levels, and attenuating iNOS and COX-2 immunoexpression (47). The antioxidant activity of MoCBP4 demonstrated in this study corroborates previous findings with a soluble lectin from *Moringa oleifera* and with seed extracts from this plant in a model of hepatic oxidative stress (48).

The cytotoxicity of MoCBP4 was evaluated by the SRB colorimetric assay in tumor cell lines (B16F10, SK-MEL, MC38) and in the normal L929 cell line at 24h and 72h. MoCBP4 did not exhibit cytotoxicity under any of the tested conditions, corroborating previous studies with other cell lines (10). This lack of cytotoxicity is a promising result, indicating that MoCBP4 should not exacerbate the side effects commonly associated with anticancer substances. Demonstrated that SN-38, the active metabolite of irinotecan, is considerably more cytotoxic than irinotecan itself in intestinal epithelial cells. SN-38 exhibits high anticancer potency, but is limited by its hydrophobicity and instability (49). Drug delivery systems, especially protein nanoparticles, seek to optimize SN-38 delivery by improving its solubility, stability, targeting, and bioavailability for clinical application (50). SN-38 conjugates with human serum albumin (HSA) have already been developed and have shown promising results as anticancer agents (51). To evaluate the possible interference of MoCBP4 with SN-38 cytotoxicity, assays were performed with MC38 cells, combining different concentrations of the protein with SN-38 (2.5 µM) for 48h. The results indicate that MoCBP4 did not alter the cytotoxic profile of SN-38 at the tested concentrations. *In vivo* assays were then conducted to investigate the effect of MoCBP4 on irinotecan chemotherapy.

The immune system plays a crucial role in cancer development and progression, with immunosuppressive cells contributing to tumorigenesis, especially in patients with immune-related disorders (52). The MC38 tumor transfer model in mice, mimicking human colon cancer (53), was used to evaluate the effect of MoCBP4 on tumor growth. All groups received 500,000 MC38 cells. Tumor growth was observed in all groups; however, growth in the irinotecan-treated groups was minimal, confirming the drug’s efficacy in this model (54). Tumor measurement after euthanasia confirmed that irinotecan reduced tumor growth threefold compared to the control group. MoCBP4 did not interfere with the antitumor effect of irinotecan. Myeloperoxidase (MPO) in the duodenal tissue was used to assess the systemic effect of treatment. As expected, irinotecan induced neutrophil migration, which was attenuated by MoCBP4. This result suggests that MoCBP4 acts on the side effects of irinotecan without affecting its antitumor activity. The reduction of neutrophil migration by MoCBP4 has been demonstrated in other inflammatory models, such as zymosan-induced peritonitis, carrageenan-induced paw edema (10), and acute pancreatitis (47), suggesting its systemic action. While effective against tumors, chemotherapeutic drugs can damage healthy tissues, causing toxic side effects. Onivyde, an irinotecan nanoliposome, offers improved pharmacokinetics and tumor biodistribution, but still causes mucositis. Berberine, a natural alkaloid, has shown beneficial effects on intestinal mucositis and anticancer synergy in combination with cytotoxic drugs (55). Our results demonstrate that MoCBP4, co-administered with irinotecan, minimizes its side effects without compromising antitumor activity, establishing itself as a promising substance to aid in chemotherapy treatment.

## Conclusions

This study demonstrates the protective effect of MoCBP4 in irinotecan-induced intestinal mucositis. MoCBP4 improved survival, reduced leukopenia and diarrhea, attenuated inflammatory mediators and cytokines, modulated oxidative stress, and enhanced antioxidant markers, resulting in improved morpho-functional and histological parameters without interfering with irinotecan’s antitumor activity. These findings highlight MoCBP4’s pharmacological potential as a promising therapeutic candidate for mitigating chemotherapy-induced gastrointestinal toxicity.

## Acknowledgments

This work was supported by research grants from Conselho Nacional de Desenvolvimento Científico e Tecnológico (CNPq) (Award Number: SPU 07971628/2020 in called SUS/PPSUS – CE 02/2020) and Coordenação de Aperfeiçoamento de Pessoal de Nível Superior (CAPES).

